# Stability of Antibody Repertoires in Astronauts During Long-Term Spaceflight

**DOI:** 10.64898/2026.09.11.750652

**Authors:** Rohit Farmer, Yoav Pinto, Sucheta Godbole, Lu Wang, Lauren McCormick, Samuel Li, Amber Alley, Noa Deutsch, Amy R. Henry, Farida Laboune, Douglass Diak, Leonid Serebryannyy, Brian Crucian, Noemia S. Lima, Daniel C. Douek

## Abstract

Long-duration spaceflight is known to disrupt several aspects of immune function yet its impact on the antiviral antibody repertoire remains unclear. We analyzed longitudinal plasma samples from 11 astronauts participating in a 6-month long spaceflight, including preflight, inflight, and postflight timepoints. Using a high-throughput serological method based on phage display of linear peptides from thousands of viruses we assessed the stability of antiviral antibodies over time. Across two phage libraries expressing different pathogen panels, we detected antibodies against *Alphainfluenzavirus, Betainfluenzavirus, Enterovirus,* and *Orthopneumovirus consistently* with the highest abundancy in all astronauts over all timepoints. Antibody levels against these infectious agents remained remarkably stable within individuals during and after spaceflight. In contrast, substantial heterogeneity was observed among astronauts, reflecting individualized immune histories and baseline antibody profiles. These results were confirmed by a different immunoassay using whole protein from the most targeted viruses. Together, these findings demonstrate that the antiviral antibody repertoire is largely resilient during prolonged spaceflight, highlighting preserved humoral memory despite potential dysregulation in cellular immune compartments.

## Introduction

Spaceflight exposes humans to a unique combination of stressors including microgravity, pressure differences, cosmic radiation and prolonged confinement, all of which induce significant physiological changes ^1,2^. As space missions continue to increase in duration and distance, it becomes even more important that we understand how these conditions impact human health. Among the physiological systems impacted, dysregulation of the immune system represents a critical concern because it has the potential to increase susceptibility of astronauts to infections during spaceflight and after returning to Earth.

Astronauts frequently exhibit altered immune function during spaceflights, with these effects persisting for months after their return to Earth. Increases in leukocyte, granulocyte and neutrophil levels during and after flight suggest heightened innate immune activation, while changes in natural killer cell levels point to altered cytotoxic function. Broad immunological changes such as elevated inflammatory cytokines including IL-1α, IL-1β, and IL-6, and shifts in innate immune cell populations have been described ^5^. In terms of adaptive immunity, T cells are the most impacted with reduction in cell function and proliferation ^4,6^. Clinically, this dysregulation is evident in the increased rates of infectious events as well as the reactivation of latent herpesviruses, indicating impaired immune responses ^7,8^.

In contrast, studies of astronauts on long-duration missions report only minimal changes in total B cell counts, and in the distribution of B cell subsets, including naïve, memory and plasma cells ^9^. This is also reflected in plasma levels of IgM and IgG, which have been reported to remain relatively constant during spaceflight. However, alterations in IgM repertoires have been observed in cosmonauts and can persist after their return to Earth ^10^. IgA levels have been shown to increase during spaceflight and return to baseline levels post-flight (Spielmann, 2018). Similarly, in animal studies, it has been shown that despite B cell numbers may fluctuate after reentry to Earth, antibody diversity is still preserved ^11,12^. Despite these insights, a major gap remains in understanding how spaceflight affects antigen-specific antibody repertoires in humans, particularly antiviral IgG, which plays a role in long term immunity.

These outcomes highlight changes in immune defense in the space environment and underscore the need to better understand the underlying mechanisms. Here we investigated whether the levels and composition of the antiviral antibody repertoire in human plasma changes due to spaceflight. Using a high-throughput serological screening approach ^13^, we analyzed plasma samples from 11 astronauts participating in a 6 months long spaceflight, collected at seven timepoints spanning preflight, inflight and postflight. This approach enabled us to assess potential alterations in the circulating antibody repertoire against viruses during long-duration space travel, providing new insight into the humoral immune system under spaceflight conditions.

## Results

### Antibodies against *Alphainfluenzavirus, Betainfluenzavirus, Enterovirus, and Orthopneumovirus* are the most abundant in plasma

To determine whether antibody profiles remained stable or changed over time during spaceflight, plasma samples from 11 astronauts were analyzed across seven longitudinal timepoints including preflight, inflight, and postflight: (T1) 180 days prior to spaceflight; (T2) 45 days prior to spaceflight; (T3) mid-mission; (T4) late-mission; (T5) on the day or day after of return to Earth (day 0 or 1); (T6) 30 days following return; (T7) and 90 days following return to Earth (**Figure 1A**).

**Figure 1:**
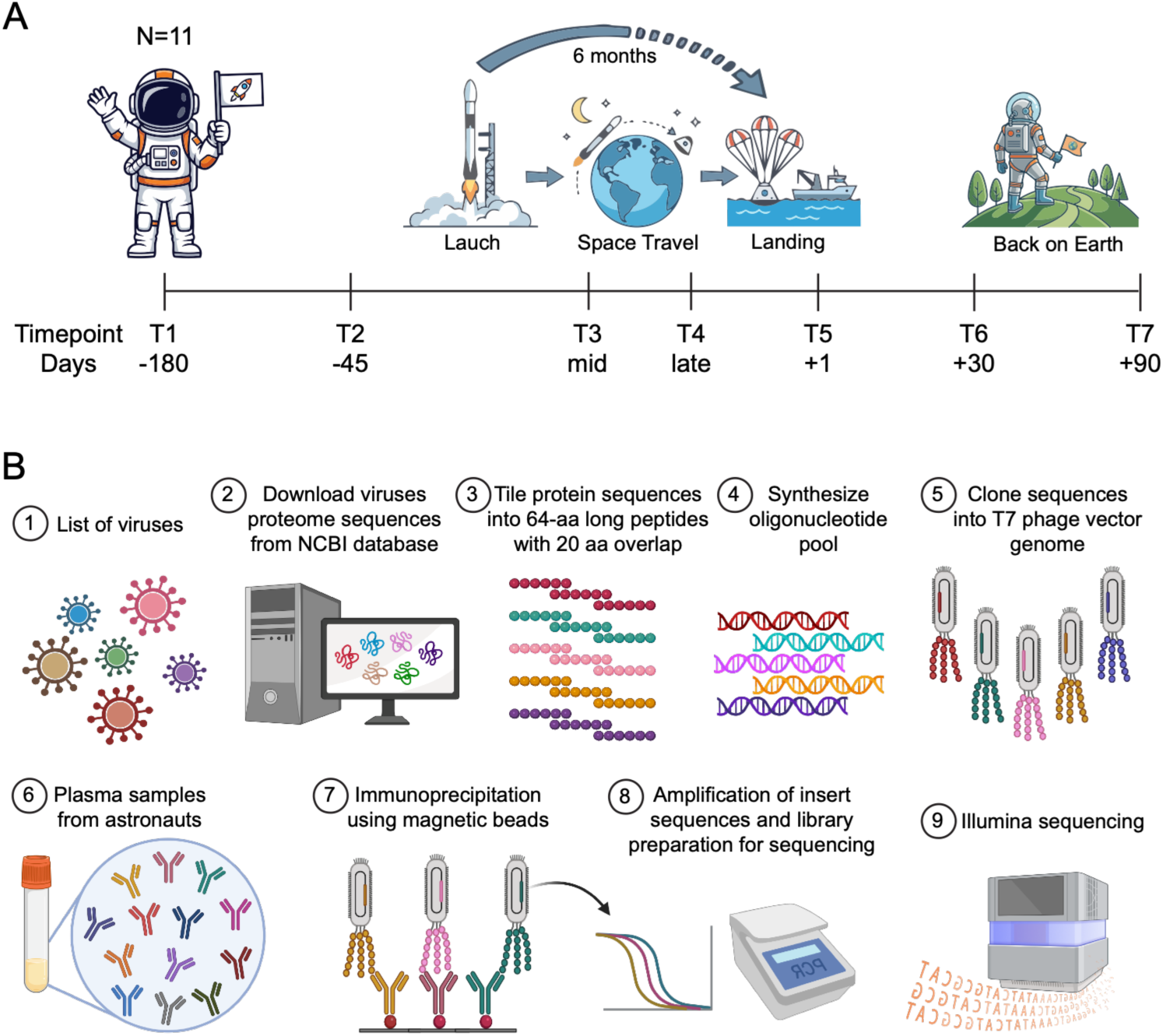
Experimental design. A) Plasma samples from 11 astronauts were analyzed across seven longitudinal timepoints: (T1) 180 days prior to spaceflight; (T2) 45 days prior to spaceflight; (T3) mid-mission; (T4) late-mission; (T5) on the day or day after of return to Earth; (TC) 30 days following return; (T7) and S0 days following return. B) PhIP-Seq overview. The top row shows the steps to create a phage library; the bottom row shows the steps to perform the immunoprecipitation and sequencing experiment. Figure B was partially created in BioRender. Santana lima, N. (202C) https://BioRender.com/mrlt48k.

Antimicrobial antibody repertoires were screened by a high-throughput serological profiling method based on phage display of linear peptides and next-generation sequencing, termed Phage Immunoprecipitation Sequencing (PhIP-Seq) (Mohan et al. 2018, Xu et al, 2015). In this approach, a phage library expressing peptides derived from microbial proteomes is co- incubated with plasma samples, allowing antibodies to bind their cognate peptide antigens. Antibody–phage complexes are subsequently immunoprecipitated, and the enriched phage populations are identified by next-generation sequencing (**Figure 1B**). This strategy enables the simultaneous detection of antibody reactivity against hundreds of thousands of microbial peptides, providing a comprehensive assessment of the circulating antibody repertoire.

We profiled antimicrobial antibody repertoire using two PhIP-Seq libraries: Pandemic Potential Viruses (PPV), containing 152,950 peptides derived from 19 families and 31 genera of viruses with pandemic potential, and VirScan (Xu et al 2015) that contains 128,257 peptides from additional human viruses, as well as bacteria and protozoa species. After sequencing immunoprecipitated phages, enrichment scores were calculated for each phage-displayed peptide based on a generalized Poisson regression model comparing their sequenced read counts in each sample to those of the input phage library. The numbers of significantly enriched peptides (*P*-value < 0.05) for each virus in each sample were expressed as log_2_-transformed peptide counts (tile counts). We found that antibodies targeting viruses in the genera *Alphainfluenzavirus, Betainfluenzavirus, Enterovirus,* and *Orthopneumovirus* were consistently detected with highly-enriched peptide counts across all astronauts (Figure 2). These findings indicate highly prevalent antibody responses to common respiratory and enteric viruses in the cohort. In contrast, several other viral species in the genera *Alphacoronavirus, Betacoronavirus, Lyssavirus* and others exhibited low or infrequent peptide enrichment consistent with more limited or heterogeneous exposure histories among astronauts. Collectively, these results show that antibody responses against respiratory viruses and enteroviruses are dominant in this cohort.

**Figure 2:**
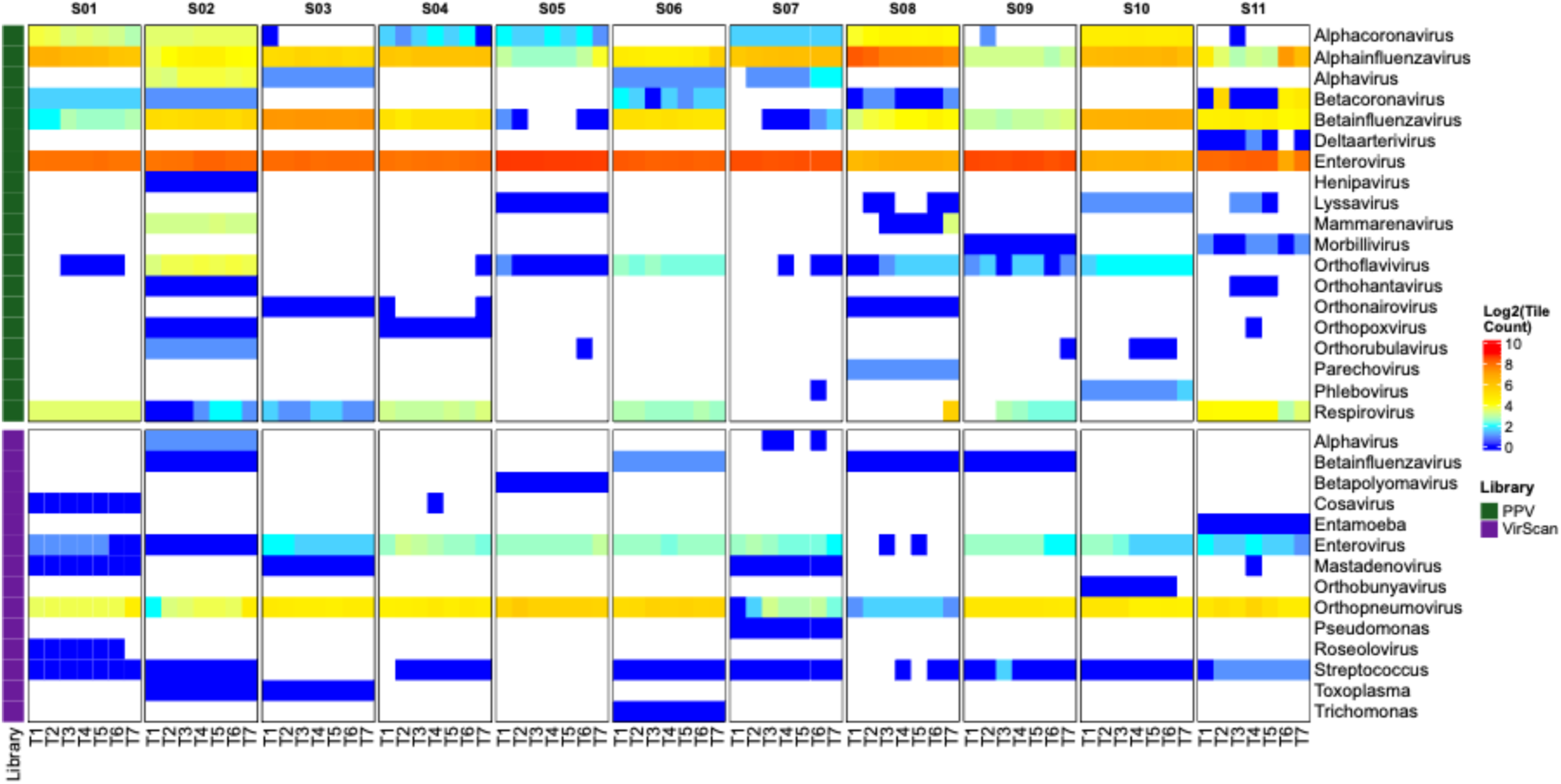
Heatmap of enriched peptide counts (log_2_-transformed) derived from the PPV library (forest green) and the VirScan library (purple). Columns represent sampling timepoints (T1-T7) grouped by astronaut (S01-S11), and rows represent viral genus for which at least one enriched peptide was detected in any astronaut at any timepoint.

### Antimicrobial antibody responses remain stable within individual astronauts during spaceflight

Longitudinal analyses of species-level antimicrobial antibody responses from the highly enriched genera in PhIP-Seq **(Supplementary Figure 5)** indicated limited variation across the seven sampling timepoints within each astronaut, consistent with the maintenance of established antibody responses during spaceflight and post-flight follow-up (**Figures 3A and B**). This pattern was particularly evident for common respiratory viruses, including *Alphainfluenzavirus influenzae (Influenza A)*, *Betainfluenzavirus influenzae (Influenza B), Enterovirus A, B, C, and D, Rhinovirus A (RVA) and C* (RVC) and *Respiratory Syncytial Virus (RSV)*.

**Figure 3:**
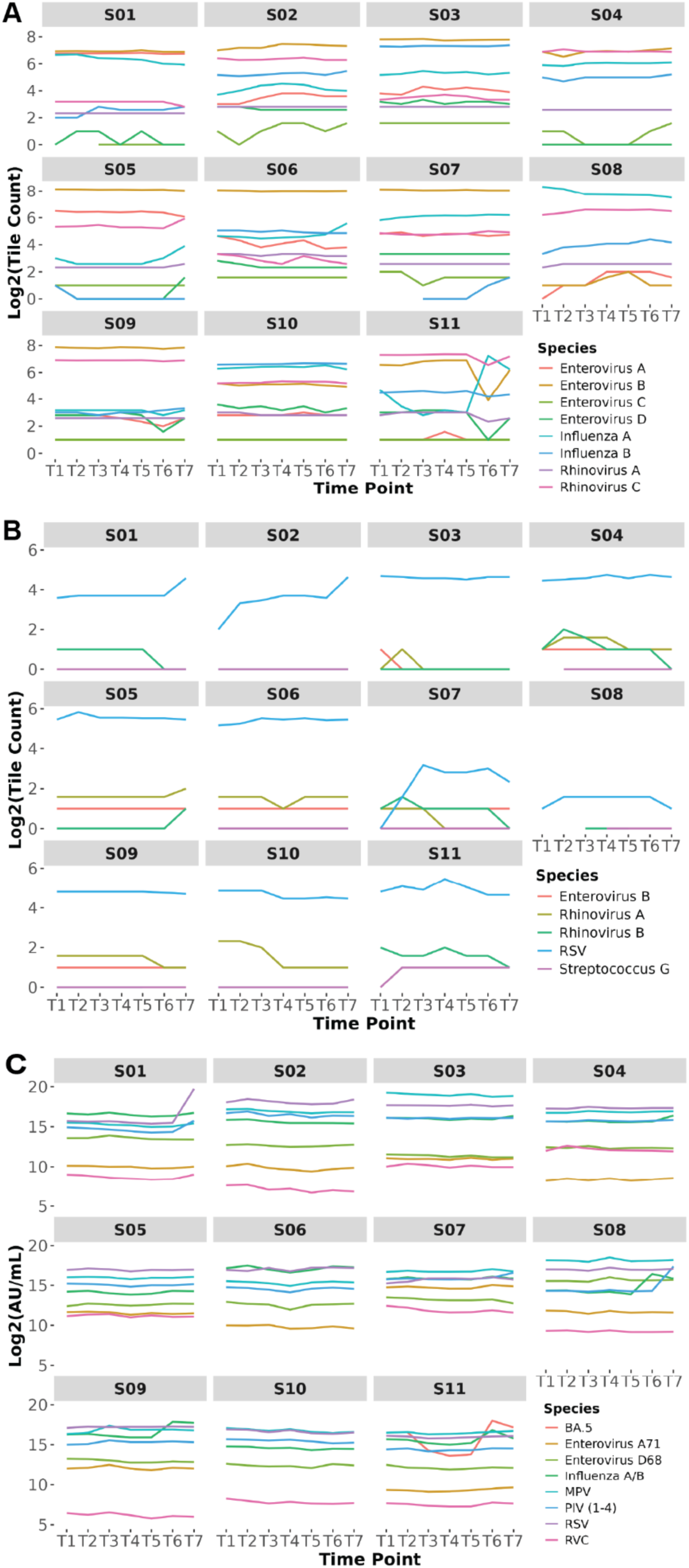
Log_2_-transformed enriched peptide count line plots derived from the PPV library (A), the VirScan library (B), and ECLIA (C). Panels A and B depict antibody responses to selected viral species over time for individual astronauts in PhIP-Seq data. Panel C depicts antibody responses to whole proteins by ECLIA over time for individual astronauts.

To formally assess how values changed over time, autocorrelation analyses were performed for each astronaut–virus pair across time lags 1–7 (T1-T7), and significance was evaluated using the Ljung–Box test **(Supplementary Figures 6A and B)**. Autocorrelation analysis examines how a variable relates to its own past values over successive time lags, helping to identify patterns over time. The Ljung–Box test is a statistical test that evaluates whether a group of autocorrelations, taken together across multiple lags, differs significantly from zero, indicating the presence of non-random structure in the data. In the PPV dataset, nominally significant time dependence (*P-*value < 0.05) was only detected for *Influenzae A* in Astronauts 1 (S01) and 8 (S08), *Influenzae B* in Astronaut 7 (S07), and *Rhinovirus C* in Astronaut 3 (S03) **(Supplementary Table 1 PPV)**. In the VirScan dataset, nominal autocorrelation was observed for *RSV* in Astronaut 2 (S02) and *Rhinovirus A* in Astronaut 10 (S10) **(Supplementary Table 1 VirScan).** However, none of these associations remained statistically significant after false discovery rate (FDR) correction, indicating that antibody trajectories within individuals did not exhibit robust or systematic temporal changes.

To corroborate these findings, we performed electrochemiluminescence immunoassay (ECLIA) using a broad panel of respiratory viruses and enteroviruses with whole protein antigens. Consistent with the PhIP-Seq findings, ECLIA results also demonstrated stable antibody levels within individual astronauts across time for all antigens tested, including *influenza virus A and B, parainfluenza viruses (PIV 1-4), enteroviruses (EV-A71, EV-D68, and RVC)* and *RSV* (**Figure 3C**). Autocorrelation analyses identified patterns over time that were statistically significant at an unadjusted *P*-value < 0.05 for select astronaut–antigen combinations, including *PIV (types 1–4)* in Astronauts 3 (S03), 4 (S04), and 10 (S10), *Enterovirus D68* in Astronaut 7 (S07) **(Supplementary Figure 6C; Supplementary Table 1 ECLIA).** Similar to the PhIP-Seq results, none of these associations remained statistically significant after FDR correction.

Although our autocorrelation analysis demonstrated the absence of a consistent pattern over time, we did observe a few point variations, especially at post-flight timepoints. For example, Astronaut S11 exhibited a spike in anti-alphainfluenza antibodies at time point T6 both in PPV PhIP-Seq and in ECLIA results. In the ECLIA results, there was also an increase in anti-SARS-CoV-2 Omicron BA.5 antibodies at the time point T6. This was likely caused by an infection with alphainfluenza virus and a coronavirus within the first month after landing. Since these samples were collected before the emergence of SARS-CoV-2, it is likely that this astronaut was exposed to a coronavirus that induced cross-reactive antibodies with SARS-CoV-2 Omicron BA.5.

### Antiviral immune responses exhibit substantial heterogeneity across astronauts

In contrast to the relative time-dependent stability observed within individual astronauts, substantial heterogeneity in antimicrobial antibody responses was evident among astronauts.

PhIP-Seq analyses revealed pronounced inter-individual differences in the magnitude and persistence of enriched peptide counts for the same microbial species, with some astronauts exhibiting strong and sustained antibody enrichment, whereas others showed lower or undetectable responses (**Figure 4A and B**). In the PPV dataset, for example, most selected species showed log₂-transformed enriched tile counts ranging from approximately 2 to 7 among astronauts, whereas *Rhinovirus A* and *Enterovirus C* displayed comparatively narrower response patterns. Similarly, in the VirScan dataset, RSV showed marked variability in enriched tile counts among astronauts. This inter-individual variability was recapitulated in the ECLIA dataset with antibody titers for the same viral antigens varying widely and distinct response profiles (**Figure 4C**).

**Figure 4:**
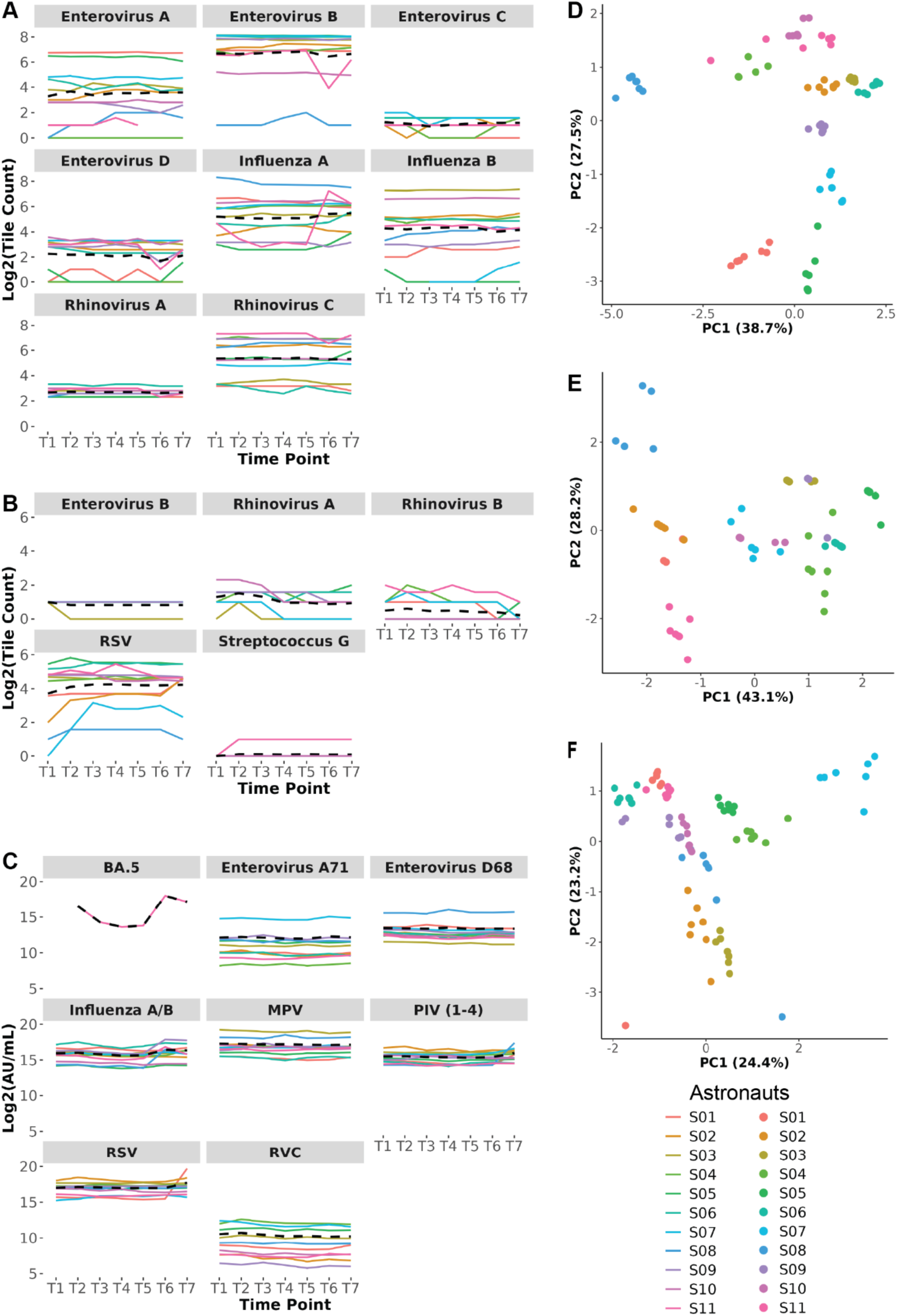
Log_2_-transformed enriched peptide counts derived from the PPV library (A), the VirScan library (B), and ECLIA (C) antibody response profiles over time for each astronaut, displayed together for the selected species and antigens. The dashed black line represents the average peptide count in A and B or antibody concentration in C across the cohort for each timepoint and antigen. Respectively, PCA plots derived from the PPV library (D), the VirScan library (E), and ECLIA (F) across all the species and antigens– each dot is a timepoint colored by astronaut.

To further explore inter-individual variation, we performed a Principal Component Analysis (PCA) in which we combined the data across all species. The separation along the principal components was largely driven by individual variability, while samples from different timepoints within the same individual remained closely grouped, showing little directional drift over time, indicating that each astronaut maintained a distinct and relatively stable antibody profile over the course of the study (**Figure 4D and E**). We observed a similar dominance of inter-individual variability across all antigens in the ECLIA data, consistent with the pattern seen in the PhIP-Seq results (**Figure 4F**).

To quantify inter-individual heterogeneity among astronauts, we fitted a series of nested linear mixed-effects regression (LMER) for each dataset. Mixed-effects models are well suited for longitudinal studies because they simultaneously account for repeated measurements within the same individual and variation between individuals, enabling separation of astronaut-specific effects from overall time dependent trends. In this framework, fixed effects estimate systematic trends shared across the cohort, such as the effect of sampling timepoint during and after spaceflight, whereas random effects account for astronaut-specific variability by allowing each astronaut to have their own baseline response level and, in some models, their own time-dependent trajectory. This approach enables separation of population-level time-dependent effects from astronaut-specific immune signatures. Four models were evaluated for each dataset: (i) an intercept-only model; (ii) a random-intercept model including astronaut identity; (iii) a model including timepoint as a fixed effect; and (iv) a model allowing astronaut-specific trajectories. Model fit was evaluated using likelihood ratio tests and Akaike Information Criterion (AIC).

Across the PPV and VirScan PhIP-Seq libraries, adding individual astronaut as a random effect produced substantial improvements in model fit relative to the intercept-only model, with ΔAIC reductions typically exceeding 150–300 units and highly significant likelihood ratio tests (FDR corrected *P*-values generally <10⁻²⁵ to <10⁻^59^), indicating strong individual effects. In contrast, adding timepoint as a fixed effect resulted in only small AIC changes (ΔAIC typically <0–10), indicating weak support for time dependent effects. Although timepoint terms were statistically significant for *Influenza A* and *Enterovirus D* in the PPV dataset and *RSV, Rhinovirus A and B,* and *Streptococcus sp. group G* in the VirScan dataset at an FDR-corrected *P*-value < 0.05, their contribution to overall variance was minimal **(Supplementary Table 2 PPV LMER, VirScan LMER)**. Allowing astronaut-specific slopes occasionally produced additional modest AIC reductions but these were substantially smaller than those attributable to the individual random intercept and did not alter the overall variance structure.

Variance decomposition confirmed that astronaut identity explained 87–99% of total variance in the PPV library and 73–99% in the VirScan panel, whereas timepoint accounted for <0.4% and 0–3%, respectively. Correspondingly, intraclass correlation coefficients (ICC) were uniformly high (0.75–0.99), demonstrating strong within-astronaut stability and pronounced inter-astronaut heterogeneity **(Supplementary Table 2 PPV ICC, VirScan ICC)**.

Applying the same modeling framework to the ECLIA assay revealed more heterogeneous variance structures across antigens. For several targets — including *Enterovirus A71*, *Enterovirus D68, MPV,* and *RVC* — the inclusion of astronaut as a random effect yielded large AIC reductions (ΔAIC ≈100–250) and highly significant likelihood ratio tests (FDR corrected *P*-values <10⁻³³ to <10⁻^49^), with ICC values of 0.94–0.98, indicating strong astronaut-level clustering. For *Influenzas A* and *B*, and *PIV*, astronaut effects remained significant but explained a smaller proportion of variance (ICC 0.53–0.65), and timepoint effects produced only modest additional improvements in fit. *RSV* exhibited comparatively weak clustering (ICC = 0.24), with most variance attributable to residual variation despite a statistically detectable astronaut effect **(Supplementary Table 2 ECLIA LMER, ECLIA ICC).**

Across all three datasets, model comparisons showed that variation in antibody signal was explained primarily by astronaut identity rather than sampling timepoint, indicating that baseline differences between astronauts dominated the antimicrobial antibody landscape. Time-dependent effects were limited, and the few nominally significant patterns did not survive multiple-testing correction, suggesting that these reflected isolated person- or antigen-specific fluctuations rather than a coordinated response to spaceflight. Thus, the concordant PhIP-Seq and ECLIA findings support a model in which humoral immune profiles remain largely stable within astronaut during the 6-month mission, while substantial inter-astronaut heterogeneity accounts for most of the observed variation.

## Discussion

Dysregulation in astronaut’s immune system during long-term spaceflight may result in higher susceptibility to infections, reactivation of latent virus, delayed wound healing, and reduced vaccine responses. In this study, we analyzed longitudinal antibody responses to microorganisms, especially viruses, from 11 astronauts across preflight, inflight and postflight to determine if antibody memory is affected by a 6-month spaceflight. We first used two PhIP-Seq libraries (PPV and VirScan) to profile antimicrobial antibody responses, then applied autocorrelation analysis to antibodies detected against selected viral species to determine whether antibody levels showed time-dependent structure consistent with immune activation, waning, or instability. ECLIA measurements were used as an orthogonal platform to compare selected antibody responses using full-length antigens. This analysis yielded four principal findings: (1) astronauts showed strong and persistent antibody recognition of several common respiratory and enteric viruses, particularly *Influenza A, Influenza B, Enterovirus A–D, Rhinovirus A and C,* and *RSV;* (2) these viruses exhibited consistently enriched peptide counts across multiple astronauts and timepoints, indicating durable antiviral antibody reactivity; (3) only a small subset of astronaut–virus combinations showed nominally significant autocorrelation at *P-*values < 0.05 in either the PhIP-Seq or ECLIA datasets, and none remained significant after FDR correction; and (4) the observed temporal variations were unique to certain astronauts and did not show a pattern across astronauts, suggesting that long-duration spaceflight was not associated with uniform or systematic waning of antiviral antibody levels.

Our results demonstrated that serological memory against historically encountered pathogens remains intact throughout spaceflight, despite well-documented evidence of T cell dysregulation, diminished NK cell function and reactivation of latent herpesviruses during flight ^4,6–8^. Our findings also highlight substantial inter-individual heterogeneity, a theme consistent with broader immunological variation reported in other populations. Such heterogeneity likely reflects differences in exposure history, vaccination status and genetic background ^14,15^. This aligns well with our finding that astronauts differed markedly in antibody magnitude despite relatively stable within-person trajectories. Vogl *et al* have shown that antibodies against gut microbiome are stable over 5 years within a person despite fluctuations in gut microbiome composition, but does not correlate between individuals ^16^. This principle was also reported in the context of self-reactive antibodies, in which IgG profiles of healthy individuals were shown to be stable over time and sufficiently individualized that longitudinal samples could be recognized by their person-specific antibody signatures ^17,18^. Our study showed that this personalized and temporally consistent feature of antibody repertoire demonstrated in the general population is applicable to astronauts, and is not affected by the spaceflight conditions.

The data should also be considered in context of the vehicle conditions onboard ISS at the time the study was performed. There clearly exists, both terrestrially and for spaceflight, a continuum of stress, immune compromise, and advent to clinical disease ^19^. Data assessing immune parameters over the lifespan of ISS has demonstrated substantial improvement in more recent crewmembers ^20^. This was attributed to the deployment of several biomedical and behavioral countermeasures to ISS, including improved resistive and aerobic exercise devices, more frequent resupply, improved communications, nutritional supplementation, and others. Evidence suggests that the improved exercise duration and loads, and maintenance of fitness in astronauts, directly correlates with reductions in latent virus reactivation ^21^. The current study was conducted during this period of ‘improved performance’ onboard the ISS, thus it is difficult to ascertain if the stabilization of humoral immunity, as presented in these results, is due to limited spaceflight induced dysregulation for these parameters per se or reflects an improvement resulting from effective countermeasures. However, the planned Artemis missions of deep space exploration will inherently amplify most mission stressors beyond those observed on the ISS, and operational constraints will simultaneously reduce the ability to implement countermeasures, logically increasing certain clinical risks.

Our study has some limitations to consider. While PhIP-Seq is a powerful high-throughput approach, it is limited by its reliance on linear peptides, which may fail to capture conformational or discontinuous epitopes present in native protein structures. In this context, the ECLIA platform provided an important complement by utilizing full-length proteins that better preserve native conformations and structural epitopes. The consistency of results across these independent and methodologically distinct platforms strengthens the conclusion that the humoral immune system remains broadly stable under long-duration spaceflight conditions, with only isolated instances of variation. The cohort size, though typical for astronaut studies, restricts statistical power and generalizability.

In summary, our study demonstrates that despite known immune dysregulation in other cellular compartments during long duration spaceflight, the antimicrobial antibody repertoire remains remarkably stable within individual astronauts which should provide protection against infection under these physiologically stressful conditions.

## Methods

### Cohort and sample collection

Eleven International Space Station astronauts participated in the ‘Functional Immune’ study which was operational from 2017-2020. Four of the astronauts were female, and seven were male. All the astronauts flew to ISS on the Russian ‘Soyuz’ vehicle, with the exception of one who flew on Space-X Falcon 9 Dragon. Mission durations for all astronauts were approximately 6 months. Institutional review board approval was obtained from the Committee for the Protection of Human Subjects at the Johnson Space Center (JSC), Houston, Texas. Informed consent was obtained from all astronauts who participated in the study.

Blood samples were collected at pre-flight, in-flight, and post-flight time points, according to the experimental design (**Figure 1A**). For in-flight time points astronauts collected blood samples from each other via standard venipuncture protocol, immediately prior to hatch closure and vehicle return to Earth. The ‘MID’ in-flight samples were returned by any returning vehicle near the middle of the crewmember’s mission. The specific timing was dictated operationally by the returning vehicle. The ‘LATE’ in-flight samples returned with the crewmember at the end of the mission.

Blood samples were collected into tubes containing ACD anti-coagulant preservative and stored in thermo-stable pouches to maintain ambient temperature. The time elapsed between needle-stick on ISS and arrival of the samples at the JSC laboratory in Houston could vary but averaged 36 hours.

To account for any differences in sample storage due to the longer transportation of in-flight samples versus those collected on Earth (pre or post-flight), all baseline samples were aged similarly before processing. The aged ACD blood samples were centrifuged at 974rcf (2200rpm) for 22 minutes to isolate plasma. Aliquots of plasma were collected and immediately frozen at -80°C. Samples remained frozen until PhIP-Seq and ECLIA processing.

### PhIP-Seq

The Pandemic Potential Viruses (PPV) phage library was designed to target viruses with known or potential outbreak risk and developed as described previously ^13,22^ and outlined in **Figure 1B**. PPV comprised 152,950 peptides derived from 19 families and 31 genera. Each peptide was 64 amino acids in length and designed with a 20–amino acid overlap with adjacent peptides originating from the same protein sequence. Redundancy within the library was minimized such that no pair of peptides shared greater than 98% sequence identity. The peptide sequences were reverse translated into DNA sequences using *E. coli* codon usage and prefix/suffix sequences were attached (prefix: CGCAAATGGGCGGTAGAATTCT; suffix: TAGAAGCTTAGGTGAGATGACAGG) to make the oligonucleotide sequences. An oligonucleotide pool (Twist Bioscience) containing all the sequences was amplified by PCR (forward primer: CGCAAATGGGCGGTA; reverse primer: CCTGTCATCTCACCTAAGC). The prefix/suffix adaptors were then excluded by a restriction enzyme digestion with the EcoR I and Hind III enzymes (New England Biolabs). The digested products were purified using SPRIselect Reagent Kit (Beckman Coulter) at 2X concentration and cloned into the T7 phage vector arms using the T7Select 10-3 Cloning Kit (Millipore Sigma). Phages were grown in *E. coli* BLT5403 on LB agar plates containing 50µg/mL ampicillin, collected in a phage extraction buffer (20 mM Tris-HCl, 100 mM NaCl, 6 mM MgSO4, pH 8.0), and frozen with 10% DMSO and 50 µg/mL of kanamycin.

The immunoprecipitation reactions were performed as described in the published protocol (Mohan et al, 2018) with a few modifications. The optimal amount of phages per sample was tested within a range of 8,000 to 200,000-fold coverage for each phage peptide sequence and defined at 40,000-fold coverage. The concentration of total IgG in each plasma sample was measured by ELISA. The plasma samples were then diluted to 200µg/mL in PBS and 10µl was mixed with 1mL of the phage library diluted in phage extraction buffer. This mixture was incubated overnight at 4°C with end-over-end rotation. For capture of human IgG and IgA, 20µL per sample of protein G-coated Dynabeads (ThermoFisher Scientific) and 20µL per sample of protein A-coated Dynabeads (ThermoFisher Scientific) were washed in PBS + 0.02% Tween-20, added to each well of the phage immunoprecipitation mixture, and incubated at 4°C for 4 hours with end-over-end rotation. The plate was placed on a magnet and supernatant was removed, followed by 5 washes with IP wash buffer (150mM NaCl, 50mM Tris-HCL, 0.1% NP-40, pH 7.5). Beads were transferred to PCR plates and frozen.

To sequence the insert of the immunoprecipitated phages, three PCR reactions were run as described previously (Mohan et al, 2018) with a few modifications. For immunoprecipitation reactions using PPV phage library, PCR1 used forward primer tcgtcggcagcgtcagatgtgtataagagacagATGCTCGGGGATCCGAATTC, and reverse primer gtctcgtgggctcggagatgtgtataagagacagCTAGTTACTCGAGTGCGGCC, and anealing temperature was changed to 60°C. For immunoprecipitation reactions using VirScan phage library, PCR1 used forward primer tcgtcggcagcgtcagatgtgtataagagacagGGTGTGATGCTCGGGGATCC, and reverse primer gtctcgtgggctcggagatgtgtataagagacagAGTTACTCGAGCTTATCGTC. PCR1 product was purified using SPRIselect Reagent Kit (Beckman Coulter) at 0.5X concentration. PCR2 used 1µL of purified PCR1 product, 2.5µL of IDT for Illumina DNA/RNA UD indexes, and anealing temperature was changed to 63°C. PCR2 products (5µL per sample) were pooled in one tube per plate and the pools were purified twice using SPRIselect Reagent Kit (Beckman Coulter) at 0.8X concentration. The DNA concentration was quantified using a Qubit dsDNA Quantification Assay Kit (ThermoFisher Scientific), and 250ng of purified PCR2 products were added to PCR3 reaction. The product of the PCR3 reaction was purified twice using SPRIselect Reagent Kit (Beckman Coulter) at 0.8X concentration, and verified by BioAnalyzer using Agilent High Sensitivity DNA Kit (Agilent). The final library was sequenced on a NextSeq 2000 using a NextSeq™ 1000/2000 P3 XLEAP-SBS™ Reagent Kit (300 Cycles).

All samples were immunoprecipitated with PPV and VirScan in separate plates with technical replicates, and sequenced at higher than 10-fold coverage (10x number of phages in the library). In each experiment, in addition to plasma samples derived from NASA astronauts, we included 2 types of controls: (1) mock samples consisting of phages without plasma; (2) no-template controls (NTCs; water). In addition, the input PPV and VirScan phage libraries were sequenced to an average depth of approximately 100-fold coverage and were not subjected to immunoprecipitation.

### ECLIA

Samples were tested across a custom 10-spot multiplex 96-well Meso Scale Discovery (MSD) electrochemiluminescence immunoassay (ECLIA) as previously described ^23^. Antigens for enterovirus-D68 (EV-D68), enterovirus-A71 (EV-A71), respiratory syncytial virus (RSV-A strain A2), human metapneumovirus (HMPV), influenza virus A/B, parainfluenza virus (PIV 1-4), SARS-CoV-2 BA.5 Spike, and Rhinovirus C (RVC) were included.

To perform the assays, plates were blocked by adding MSD Blocker A and incubating for 1 hour. The plates were washed and 25µL of diluted sample or reference standard was added to each well and incubated with shaking. Samples were diluted 1:1000 followed by 3, 10-fold serial dilutions. The plates were washed, 25µL of 1X Sulfo-Tag labelled anti-human IgG detection antibody was added (MSD, D21ADF-3) to each well, and plates were incubated with shaking for 1 hour. The plates were washed and 1X MSD Gold Read Buffer B was added. Plates were analyzed using the MSD Sector Imager S600 to generate ECL signals for each array element. The signals from the reference standard as a function of concentration (in arbitrary units per mL; AU/mL) were fit to a four-parameter logistic curve. Sample IgG binding was interpolated to the reference standard curve and average dilution-adjusted concentrations were reported.

### Computational analysis

#### PhIP-Seq

Our PhIP-Seq analysis workflow comprised two major steps. In the first step, enrichment scores were assigned to each peptide tile. Analysis began with demultiplexed sequencing reads ^24^ followed by library-specific read trimming ^25^ and alignment to a reference Bowtie index to obtain read counts ^26^. Samples with low read depth were excluded, after which counts were normalized and enrichment scores were calculated as −log_10_(*P*-values) derived from a generalized Poisson regression comparing sample read counts to those of the input phage library ^13^.

In the second step, quality control (QC) was performed by assessing the distributions of read counts across all sample types, including the phage libraries **(Supplementary Figure 1)**, and by evaluating correlations of normalized read counts across technical replicates and sample types. For all the samples, replicates correlated well (r >= 0.9) **(Supplementary Figure 2 and 3)**, enrichment scores were averaged across replicates. A mock sample–based threshold was then calculated to define a background cutoff, enabling exclusion of peptides that did not exceed background signal as determined by the mock-sample score distribution **(Supplementary Figure 4)**. Peptides that passed this threshold were considered enriched and subsequently used for downstream visualization and statistical analysis **(Supplementary Figure 5**).

#### Autocorrelation Analysis

Autocorrelation analysis was performed to evaluate the time dependent structure of antibody responses measured at seven time points. This analysis was conducted independently for each astronaut and for each species of interest in both PhIP-Seq libraries—PPV and VirScan—and subsequently for multiplex immunoassay data from ECLIA.

For the PhIP-Seq datasets, for each astronaut–species pair, the log₂-transformed enriched peptide counts across the seven ordered timepoints were used to compute the autocorrelation function (ACF). The ACF was estimated for lags 1 through 7 using the standard definition of the sample autocorrelation coefficient implemented in R. The Ljung– Box test was applied to assess whether the observed autocorrelation structure differed significantly from that of white noise. The resulting *P*-values were corrected for multiple testing using a false discovery rate (FDR) adjustment. Species for which the test reached statistical significance at FDR corrected *P-value* < 0.05 were flagged as having non-random time dependence.

A parallel autocorrelation analysis was performed for ECLIA immunoassays. For each astronaut and each analyte, the measured AU/mL values across the seven time points were used as input to compute ACF values and corresponding Ljung–Box statistics using the same analytical framework applied to the PhIP-Seq data.

#### Linear Mixed-Effects Regression and Variance Decomposition

Linear mixed-effects regression (LMER) was performed to quantify inter-individual heterogeneity and time dependent variation in antibody responses measured at seven timepoints. Analyses were conducted independently for each species of interest within both PhIP-Seq libraries (PPV and VirScan) and for each antigen measured on the ECLIA multiplex immunoassay platform. This modeling framework enabled direct comparison of inter-individual and time dependent contributions to signal variability across the PPV, VirScan, and ECLIA datasets.

For each dataset, a series of nested linear mixed-effects models was fit using the lme4 package in R ^27^. Four model structures were evaluated: (i) an intercept-only model (tile_count ∼ 1 or ECLIA ∼ 1), (ii) a random-intercept model including astronaut identity (∼ 1 + (1 | astronaut)), (iii) a model including timepoint as a fixed effect (∼ timepoint + (1 | astronaut)), and (iv) a model allowing astronaut -specific temporal trajectories (∼ timepoint + (timepoint | astronaut)). Models were fit using maximum likelihood for model comparison and restricted maximum likelihood for final variance estimation. Model fit was assessed using likelihood ratio tests and Akaike Information Criterion (AIC), with ΔAIC values used to evaluate relative support for competing models.

To quantify the degree of within-astronaut clustering, intraclass correlation coefficients (ICCs) were computed from the variance components of the random-intercept models, including timepoint as a fixed effect, using the performance package ^28^ in R. The ICC was defined as the ratio of between-astronaut variance to the sum of between-astronaut and residual variance.

Variance decomposition was further performed to estimate the proportional contribution of astronaut identity, timepoint, and residual error to total variance. The variance attributable to timepoint was calculated as the variance of the fixed-effect predictions (Xβ), and total variance was defined as the sum of astronaut, timepoint, and residual components. Percent variance explained by each component was computed by dividing each component by total variance.

#### Principal Component Analysis

Principal component analysis (PCA) was performed to visualize global patterns of variation and assess clustering of samples (astronaut – timepoint) across species. For the PhIP-Seq datasets, matrices of enriched peptide counts were first log₂-transformed after adding a pseudo count of 1. For ECLIA, measured antibody concentrations were used as input values. For each dataset, samples were arranged as observations and viral species or antigen targets as variables, and features were centered and scaled. PCA was computed using singular value decomposition as implemented in R (prcomp). The proportion of variance explained by each principal component was calculated from the eigenvalues of the covariance matrix. Scatterplots of the first two principal components were generated to visualize clustering patterns, with each dot representing a timepoint and colored by astronaut identity. These analyses enabled unsupervised assessment of whether samples grouped primarily by individual astronauts or by timepoint across species.

## Data Availability

All the data related to the PhIP-Seq and ECLIA experiments, intermediate analysis files, and scripts are available at https://zenodo.org/records/21812100.

## Acknowledgements

We thank Dr. Thomas Vogl for helping to establish the PhIP-Seq method in our laboratory. We also thank Rahul Subramanian for the productive discussions and suggestions about the models used in this work for statistical analysis. We thank the volunteers that donated blood for this study and the spaceship crew that contributed to the sample collection and transportation. This research was supported by the Intramural Research Program of the National Institutes of Health (NIH) and by the NASA Human Research Program. The contributions of the NIH author(s) are considered Works of the United States Government. The findings and conclusions presented in this paper are those of the author(s) and do not necessarily reflect the views of the NIH or the U.S. Department of Health and Human Services.

## Author contributions

RF performed bioinformatics analysis and wrote the manuscript; YP conceptualized the project and performed PhIP-Seq experiments; SG performed bioinformatics analysis; Lu Wang performed ECLIA experiments; LM and SL generated the phage libraries used in the PhIP-Seq experiments; AA generated the sequences used in the phage library creation; ND wrote parts of the manuscript; ARH and FL performed sequencing; DD and BC conceptualized and coordinated the spaceflight study, sample collection, sample archive and sample shipment; LS coordinated ECLIA experiments; NSL coordinated PhIP-Seq experiments and oversaw manuscript preparation; DCD oversaw project conceptualization, execution and manuscript preparation.

## References

1. Tomsia, M. et al. Long-term space missions’ effects on the human organism: what we do know and what requires further research. Front. Physiol. 15, 1284644 (2024).

2. Krittanawong, C. et al. Human Health during Space Travel: State-of-the-Art Review. Cells 12, 40 (2022).

3. Crucian, B. E. et al. Immune System Dysregulation During Spaceflight: Potential Countermeasures for Deep Space Exploration Missions. Front. Immunol. 9, 1437 (2018).

4. Crucian, B. et al. Alterations in adaptive immunity persist during long-duration spaceflight. Npj Microgravity 1, 15013 (2015).

5. Marchal, S. et al. Challenges for the human immune system after leaving Earth. NPJ Microgravity 10, 106 (2024).

6. Dhar, S., Kaeley, D. K., Kanan, M. J. & Yildirim-Ayan, E. Mechano-Immunomodulation in Space: Mechanisms Involving Microgravity-Induced Changes in T Cells. Life 11, 1043 (2021).

7. Crucian, B. et al. Incidence of clinical symptoms during long-duration orbital spaceflight. Int. J. Gen. Med. 9, 383–391 (2016).

8. Rooney, B. V., Crucian, B. E., Pierson, D. L., Laudenslager, M. L. & Mehta, S. K. Herpes Virus Reactivation in Astronauts During Spaceflight and Its Application on Earth. Front. Microbiol. 10, 16 (2019).

9. Spielmann, G. et al. B cell homeostasis is maintained during long-duration spaceflight. J. Appl. Physiol. Bethesda Md 1985 126, 469–476 (2019).

10. Buchheim, J.-I. et al. Plasticity of the human IgM repertoire in response to long-term spaceflight. FASEB J. Off. Publ. Fed. Am. Soc. Exp. Biol. 34, 16144–16162 (2020).

11. Tascher, G. et al. Analysis of femurs from mice embarked on board BION-M1 biosatellite reveals a decrease in immune cell development, including B cells, after 1 wk of recovery on Earth. FASEB J. Off. Publ. Fed. Am. Soc. Exp. Biol. 33, 3772–3783 (2019).

12. Ward, C. et al. Effects of spaceflight on the immunoglobulin repertoire of unimmunized C57BL/6 mice. Life Sci. Space Res. 16, 63–75 (2018).

13. Mohan, D. et al. PhIP-Seq characterization of serum antibodies using oligonucleotide-encoded peptidomes. Nat. Protoc. 13, 1958–1978 (2018).

14. Andreu-Sánchez, S. et al. Phage display sequencing reveals that genetic, environmental, and intrinsic factors influence variation of human antibody epitope repertoire. Immunity 56, 1376–1392.e8 (2023).

15. Olin, A. et al. Demographic and genetic factors shape the epitope specificity of the human antibody repertoire against viruses. Nat. Immunol. 27, 600–612 (2026).

16. Vogl, T. et al. Population-wide diversity and stability of serum antibody epitope repertoires against human microbiota. Nat. Med. 27, 1442–1450 (2021).

17. Neiman, M. et al. Individual and stable autoantibody repertoires in healthy individuals. Autoimmunity 52, 1–11 (2019).

18. Bodansky, A. et al. Unveiling the proteome-wide autoreactome enables enhanced evaluation of emerging CAR T cell therapies in autoimmunity. J. Clin. Invest. 134, e180012 (2024).

19. Krieger, S. S. et al. Alterations in Saliva and Plasma Cytokine Concentrations During Long-Duration Spaceflight. Front. Immunol. 12, 725748 (2021).

20. Crucian, B. E. et al. Countermeasures-based Improvements in Stress, Immune System Dysregulation and Latent Herpesvirus Reactivation onboard the International Space Station - Relevance for Deep Space Missions and Terrestrial Medicine. Neurosci. Biobehav. Rev. 115, 68–76 (2020).

21. Agha, N. H. et al. Exercise as a countermeasure for latent viral reactivation during long duration space flight. FASEB J. Off. Publ. Fed. Am. Soc. Exp. Biol. 34, 2869–2881 (2020).

22. Xu, G. J. et al. Viral immunology. Comprehensive serological profiling of human populations using a synthetic human virome. Science 348, aaa0698 (2015).

23. Nguyen-Tran, H. et al. Dynamics of endemic virus re-emergence in children in the USA following the COVID-19 pandemic (2022-23): a prospective, multicentre, longitudinal, immunoepidemiological surveillance study. Lancet Infect. Dis. 26, 22–33 (2026).

24. Kang, H. M. et al. Multiplexed droplet single-cell RNA-sequencing using natural genetic variation. Nat. Biotechnol. 36, 89–94 (2018).

25. Bolger, A. M., Lohse, M. & Usadel, B. Trimmomatic: a flexible trimmer for Illumina sequence data. Bioinforma. Oxf. Engl. 30, 2114–2120 (2014).

26. Langmead, B. & Salzberg, S. L. Fast gapped-read alignment with Bowtie 2. Nat.Methods 9, 357–359 (2012).

27. Bates, D., Mächler, M., Bolker, B. & Walker, S. Fitting Linear Mixed-Effects Models Using **lme4**. J. Stat. Softw. 67, (2015).

28. Lüdecke, D., Ben-Shachar, M., Patil, I., Waggoner, P. & Makowski, D. performance: An R Package for Assessment, Comparison and Testing of Statistical Models. J. Open Source Softw. 6, 3139 (2021).

