## Supplementary Material for "Stability of Antibody Repertoires in Astronauts During Long-Term Spaceflight"

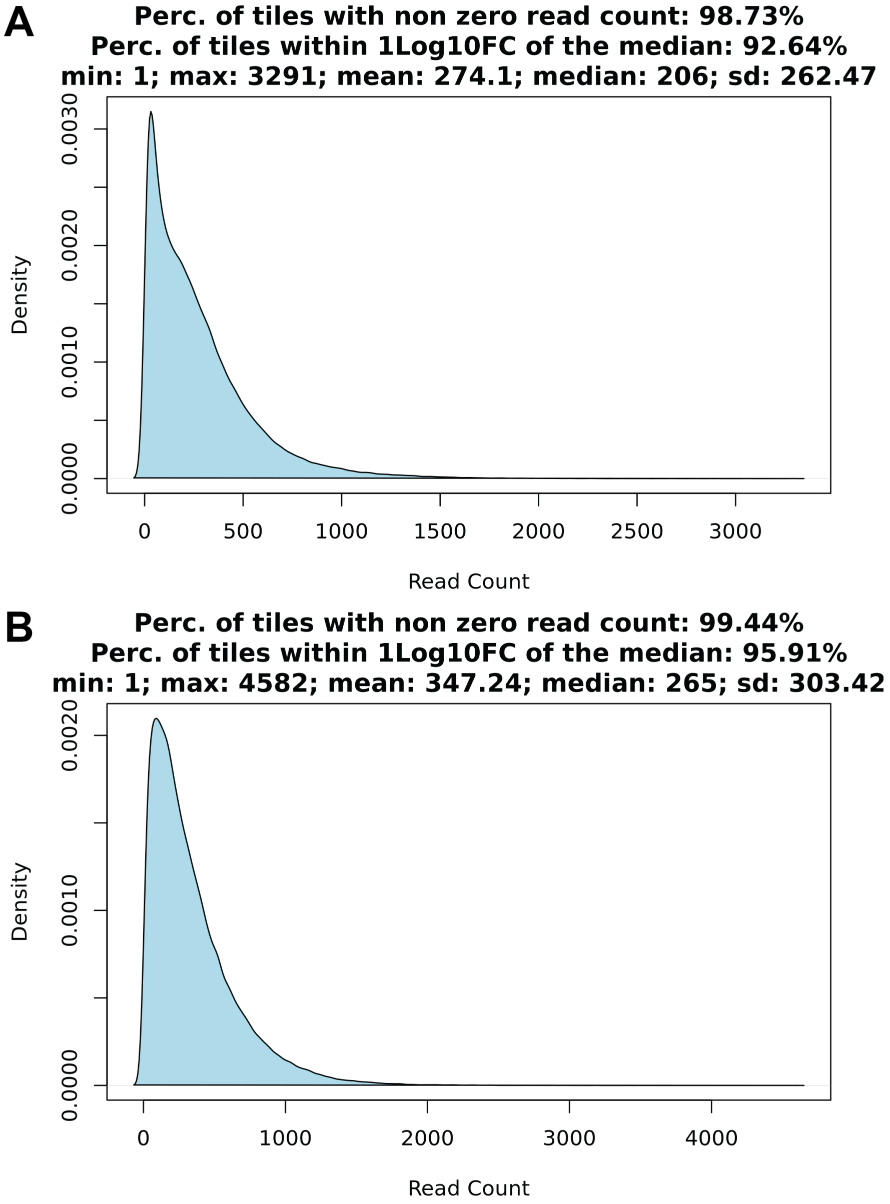


**Supplementary Figure 1.** Read count density plots for the PPV (A) and VirScan (B) libraries.

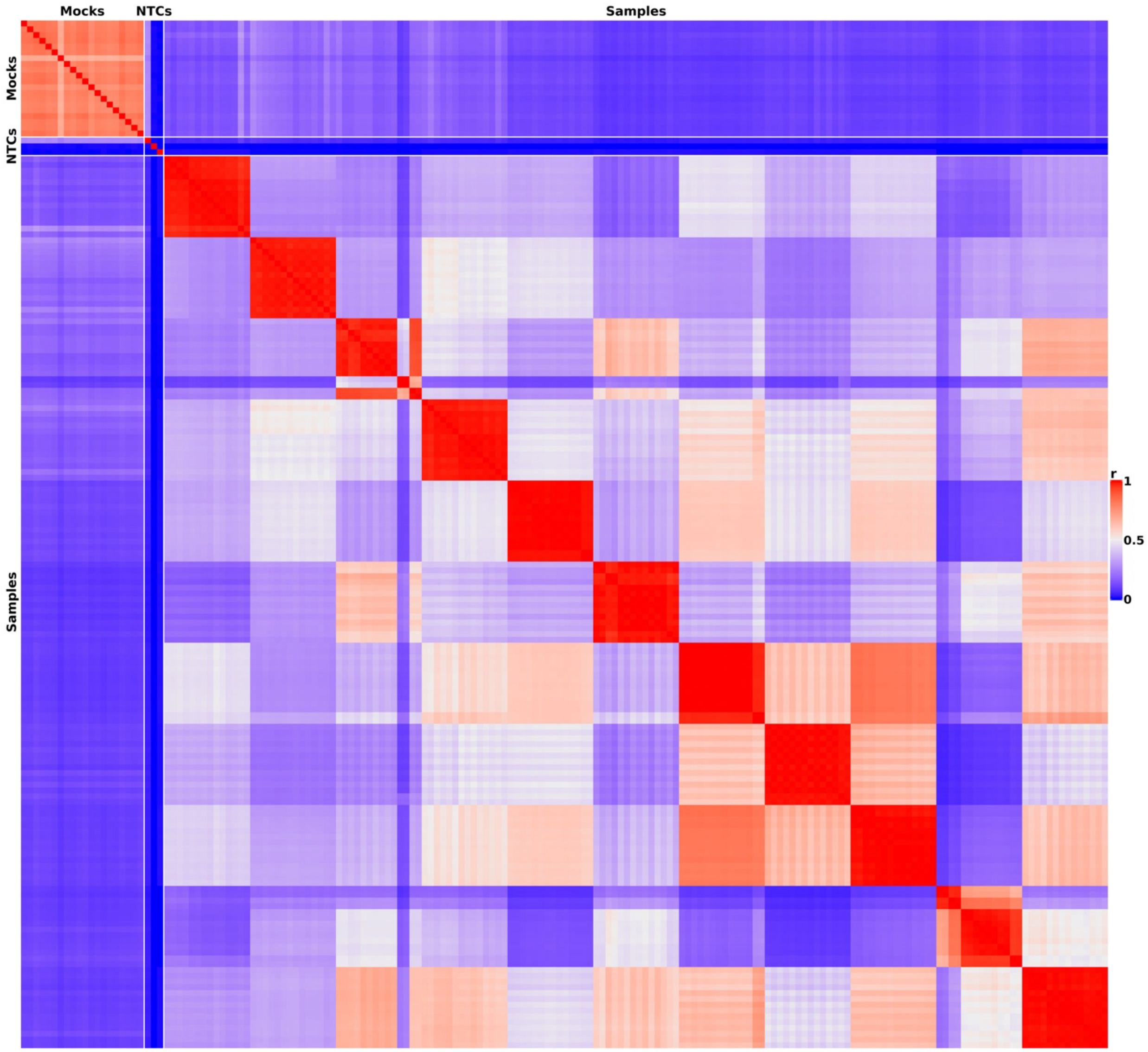


**Supplementary Figure 2.** Pairwise correlations of normalized read counts across all sample types for the PPV library. Samples are grouped by sample type along both axes, and correlations were computed using normalized read counts for all peptides. The cells display Pearson correlation coefficients, with values ranging from 0 to 1, and are color-coded on a blue-to-red scale to indicate increasing correlation strength.

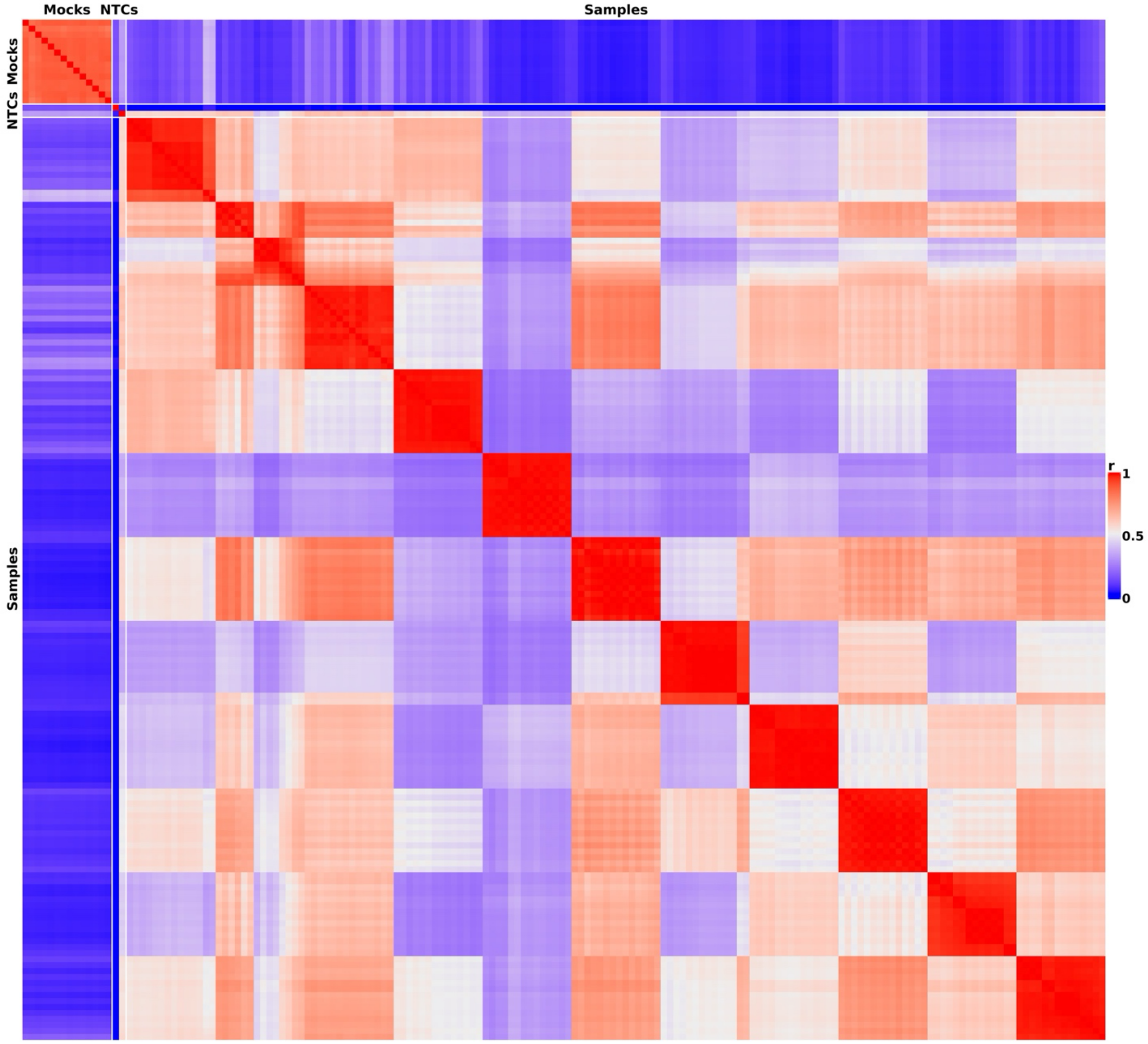


**Supplementary Figure 3.** Pairwise correlations of normalized read counts across all sample types for the VirScan library. Samples are grouped by sample type along both axes, and correlations were computed using normalized read counts for all peptides. The cells display Pearson correlation coefficients, with values ranging from 0 to 1, and are color-coded on a blue-to-red scale to indicate increasing correlation strength.

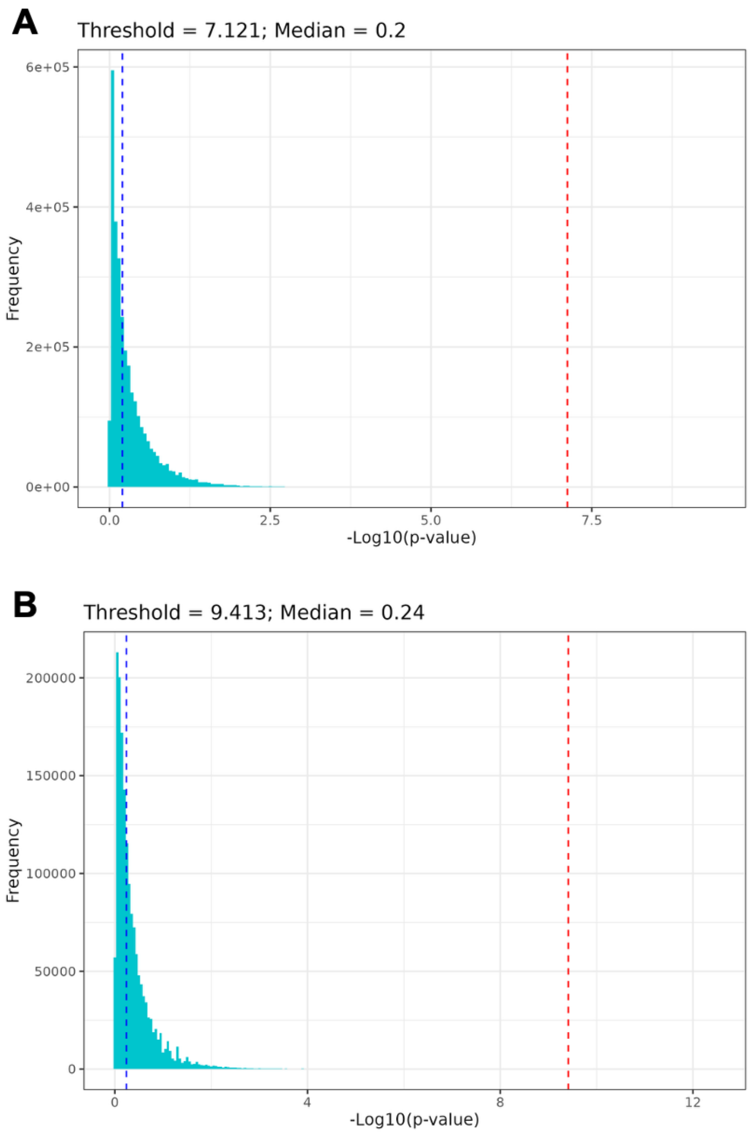


**Supplementary Figure 4.** Distribution of enrichment scores ($-\log_{10}$ p-values) for mock samples used to determine the background threshold cutoff for PPV (A) and VirScan (B) libraries. The red dashed line indicates the selected threshold, while the blue dashed line denotes the median of the distribution.

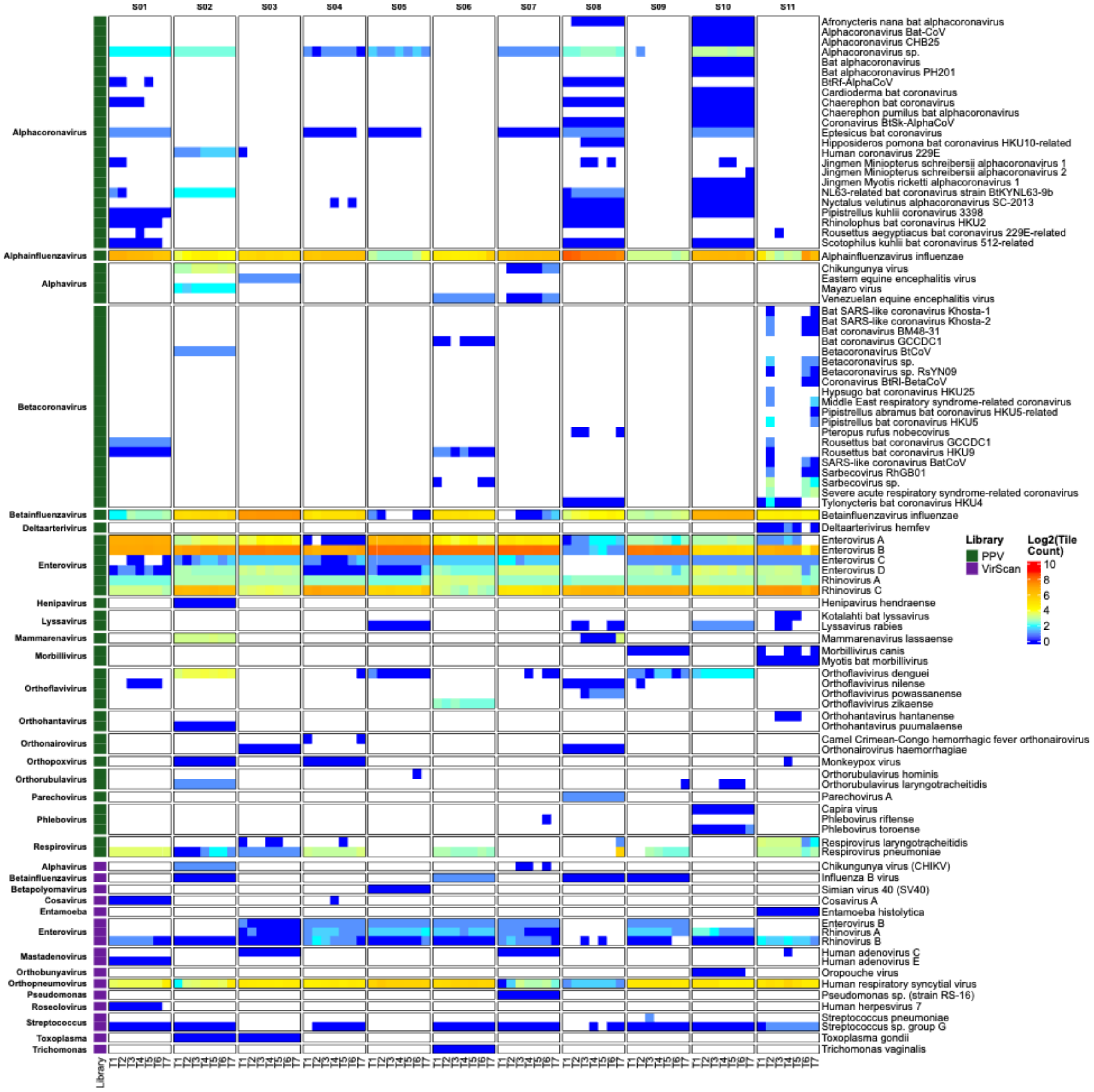


**Supplementary Figure 5.** Log2-transformed enriched peptide counts heatmaps derived from (forest green) the PPV library and (purple) the VirScan library. Columns represent sampling timepoints grouped by astronaut, and rows represent viral species for which at least one enriched peptide was detected in any astronaut at any timepoint.

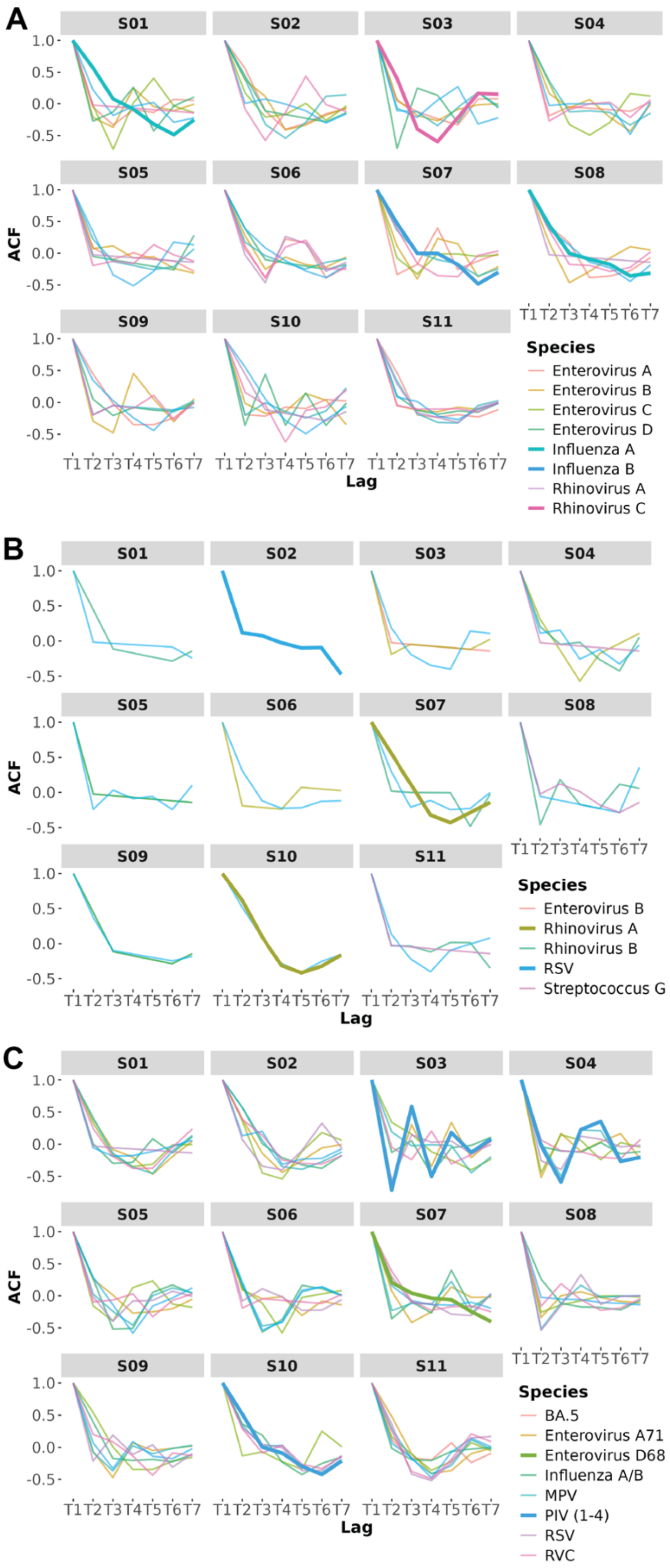


**Supplementary Figure 6:** Autocorrelation function line plots derived from the PPV library (A), the VirScan library (B), and ECLIA (C). A and B show the autocorrelation values at lags

1–7 (T1-T7) for each species per astronaut, based on the PhIP-Seq data. Panel C shows the autocorrelation values at lags 1–7 (T1-T7) for each species per astronaut, based on the ECLIA data. Bold lines indicate species for which the Ljung–Box test detected significant autocorrelation ($p < 0.05$), suggesting non-random time-dependent structure in their $Log_2$-transformed enriched peptide count trajectories. All other species, shown with standard line weight, did not exhibit statistically significant autocorrelation under the same criterion.
